# Effects of linked selective sweeps on demographic inference and model selection

**DOI:** 10.1101/047019

**Authors:** Daniel R. Schrider, Alexander G. Shanku, Andrew D. Kern

## Abstract

The availability of large-scale population genomic sequence data has resulted in an explosion in efforts to infer the demographic histories of natural populations across a broad range of organisms. As demographic events alter coalescent genealogies they leave detectable signatures in patterns of genetic variation within and between populations. Accordingly, a variety of approaches have been designed to leverage population genetic data to uncover the footprints of demographic change in the genome. The vast majority of these methods make the simplifying assumption that the measures of genetic variation used as their input are unaffected by natural selection. However, natural selection can dramatically skew patterns of variation not only at selected sites, but at linked, neutral loci as well. Here we assess the impact of recent positive selection on demographic inference by characterizing the performance of three popular methods through extensive simulation of datasets with varying numbers of linked selective sweeps. In particular, we examined three different demographic models relevant to a number of species, finding that positive selection can bias parameter estimates of each of these models—often severely. Moreover, we find that selection can lead to incorrect inferences of population size changes when none have occurred. We argue that the amount of recent positive selection required to skew inferences may often be acting in natural populations. These results suggest that demographic studies conducted in many species to date may have exaggerated the extent and frequency of population size changes.

## INTRODUCTION

The widespread availability of population genomic data has spurred a new generation of studies aimed at understanding the histories of natural populations from a host of model and non-model organisms alike. In particular, genome-scale variation data allows for inference of demographic factors such as population size changes, the timing and ordering of population splits, migration rates between populations, and the founding of admixed populations (Pool *et al.* 2010; Pickrell and Pritchard 2012; Sousa and Hey 2013). Such efforts can refine our picture of demographic events inferred from the archaeological record (e.g. Fagundes *et al.* 2007; Goebel *et al.* 2008), or reveal such events in species where no archaeological data are available, and can aid conservation efforts by complementing census data (e.g. Hájková *et al.* 2007; Garrick *et al.* 2015).

Population genomic datasets are well suited for this task simply because demographic changes leave their mark on patterns of genetic variation. Recent population growth for example will result in an excess of rare variation compared to equilibrium expectations (Fu 1997), while population contraction will result in an excess of intermediate frequency alleles (Maruyama and Fuerst 1985). In recent years researchers have devised a variety of methods that seek to detect the population genetic signatures of these demographic events. These include approximate Bayesian computation (ABC) methods, where simulation is used to approximate the posterior probability distributions of a demographic model’s parameters through the use of a collection of population genetic summary statistics without specification of an explicit likelihood function (Tavaré *et al.* 1997; Pritchard *et al.* 1999; Beaumont *et al.* 2002; Marjoram *et al.* 2003; Excoffier *et al.* 2005; Wegmann *et al.* 2010). Other approaches, such as ∂a∂i (Diffusion Approximations for Demographic Inference; Gutenkunst *et al.* 2009), use the probability density of the site frequency spectrum (SFS) under a given demographic model and parameterization to calculate the likelihood of the observed SFS (Marth *et al.* 2004; Gutenkunst *et al.* 2009), thereby allowing for optimization of model parameters. More recently, methods based on the sequentially Markovian coalescent (SMC; McVean and Cardin 2005; Marjoram and Wall 2006) have been devised (Li and Durbin 2011; Sheehan *et al.* 2013; Schiffels and Durbin 2014), to infer how a population’s size has changed over time through the description of patterns of genetic variation along a recombining chromosome.

Applications of these inference methods have revealed much about the demographic histories of various species. For instance, early studies of human genomic variation found that non-African populations experienced a considerable population bottleneck (Marth *et al.* 2003), most likely associated with migration out of Africa (Reich *et al.* 2001; Adams and Hudson 2004; Voight *et al.* 2005), followed by more recent recovery. Later studies refined the estimated timing of this bottleneck to ∼50 kya (Fagundes *et al.* 2007; Gravel *et al.* 2011; Lukić and Hey 2012), uncovered a second bottleneck associated with the divergence of European and Asian populations (Keinan *et al.* 2007; Gutenkunst *et al.* 2009; Gravel *et al.* 2011), and inferred that the recovery from this bottleneck proceeded through continuous exponential growth (Fagundes *et al.* 2007; Gutenkunst *et al.* 2009). Numerous studies have found strong genetic signals of admixture in many different human subpopulations (Meinilä *et al.* 2001; Parra *et al.* 2001; Martínez-Cortés *et al.* 2012; Patterson *et al.* 2012; Moorjani *et al.* 2013; Auton *et al.* 2015). Finally, recent studies of large population samples capable of observing very rare alleles found evidence that this growth has accelerated dramatically within the last several thousand years (Coventry *et al.* 2010; Tennessen *et al.* 2012; Gao and Keinan 2016). Similarly, recent studies in *Drosophila melanogaster* show evidence of a severe out-of-Africa bottleneck (Begun and Aquadro 1993), but occurring within the last 20,000 years (Li and Stephan 2006; Thornton and Andolfatto 2006). There is also growing support from population genetic studies for African-European admixture in the North American population of *D. melanogaster* (Duchen *et al.* 2013; Bergland *et al.* 2015; Kao *et al.* 2015).

While demographic inference from population genomic data in its various forms has proven to be successful technique, a unifying assumption of these various inference methods (ABC, SFS-based, and SMC-based approaches.) is that the genetic data in question are strictly neutral and free from the effects of linked selection in the genome. While this is an important simplifying assumption, it may be the case that in many populations a sizeable fraction of the genome is influenced by natural selection (Hahn 2008; Sella *et al.* 2009; Corbett-Detig *et al.* 2015). Indeed natural selection can produce skews in patterns of genetic variation that are quite similar to those generated by certain non-equilibrium demographic histories. For example, positive selection driving a mutation to fixation (i.e. a selective sweep; Maynard Smith and Haigh 1974) may resemble a population bottleneck (Simonsen *et al.* 1995). Conversely, many demographic perturbations are well known to cause unacceptably high rates of false positives for many classical tests for selection (Simonsen *et al.* 1995; Przeworski 2002; Akey *et al.* 2004; Jensen *et al.* 2005; Nielsen *et al.* 2005). Thus, if natural selection has a substantial impact on genome-wide patterns of variation, then many demographic parameter estimates could be biased (Hahn 2008; Gazave *et al.* 2014). Indeed this has been shown to be the case for at least some scenarios of background selection (Ewing and Jensen 2016), where purifying selection reduces levels of neutral polymorphism at linked sites.

Here, we examine the potential impact of linked positive selection on three of the most widely used methods for demographic inference: ABC (Pritchard *et al.* 1999; Beaumont *et al.* 2002), ∂a∂i (Gutenkunst *et al.* 2009), and PSMC (pairwise sequentially Markovian Coalescent; Li and Durbin 2011). We demonstrate that selection can substantially bias parameter estimates, often leading to overestimates of the severity of population bottlenecks and/or the rate of population growth. Moreover, we show that selective sweeps can result in the selection of the incorrect demographic model: if a reasonably small fraction of loci used for inference are linked to a selective sweep, one may incorrectly infer that a constant-size population experienced a bottleneck. Finally, we discuss the implications of our results for inferences made in humans and *Drosophila*, and recommend steps that could partially mitigate the bias caused by selection.

## RESULTS

### The impact of positive selection on variation under four demographic scenarios

In order to test the potential impact of positive selection on demographic inference, we simulated four different population size histories: a constant population size model, a bottleneck model, an exponential growth model, and a population contraction followed by later exponential growth (Figure 1; Methods). We begin by demonstrating the impact of linked selective sweeps on genetic diversity, as measured by *π* (Nei and Li 1979) and Tajima’s *D* (Tajima 1989), under each of these models. For each model, the selection coefficient, *s*, for each sweep was set to 0.05, and selection was modeled as a “hard” sweep involving the single origin of an advantageous mutation going to fixation (Methods). In Figure 1, we show the mean values of these two statistics at increasing distances from a selective sweep (where distance is measured by the total crossover rate, *c*, over *s*) under each of the four demographic models examined.

**Fig. 1.**
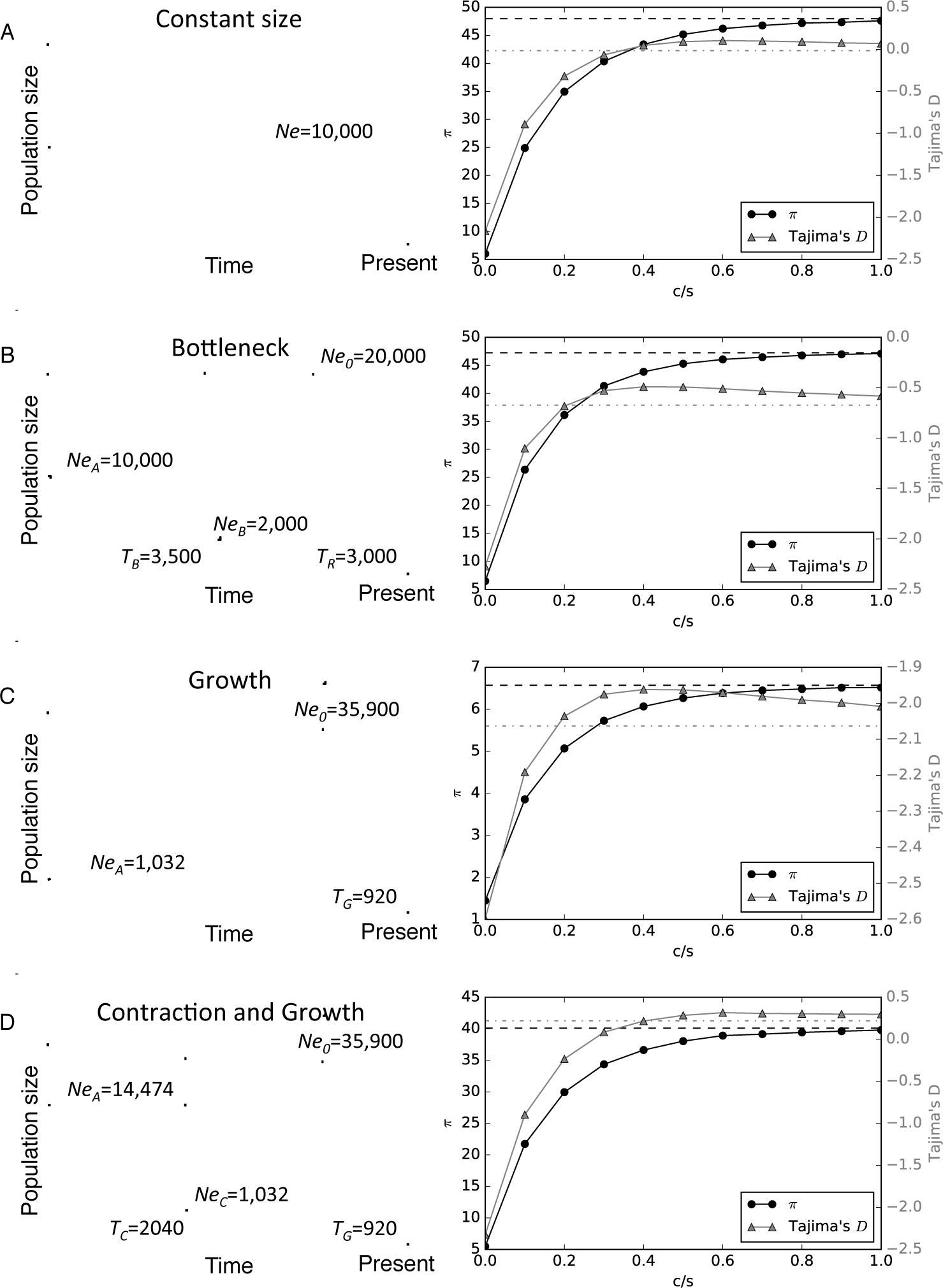
Demographic models used in this study. For each model, a diagram of the population size history is shown on the left (not to scale) along with the values of each parameter. On the right, the values of *π* and Tajima’s *D* are shown for windows sampled at varying distances (measured by the total crossover rate over the selection coefficient, *c*/*s*) from a hard selective sweep. (*A*) A model with constant population size. (*B*) A population bottleneck (parameterization from Marth *et al.* 2004). (*C*) Recent exponential population growth. (*D*) A three-epoch model with a population contraction and recent exponential growth (a simplified version of the European model from Gravel *et al.* 2011). Dotted lines in each right panel indicate the means of *π* (black dashes) and Tajima’s *D* (gray dash-dots) respectively from neutral coalescent simulations.

Under our constant population size model, a recent selective sweep with *s*=0.05 has a marked effect on genetic polymorphism at the site of the sweep (Figure 1A): on average *π* is reduced approximately 8-fold (6.01, versus a neutral expectation of 48.0), and Tajima’s *D* is well below zero (mean *D*: -2.16). At increasing distances, both of these statistics recover toward their expectation under neutrality, which they have nearly reached at *c*/*s*=1. At intermediate distances Tajima’s *D* passes through a range where its value is above the expectation of approximately zero, as has been observed previously (Teshima *et al.* 2006; Schrider *et al.* 2015). Under our bottleneck scenario (Figure 1B), which is Marth et al.’s (2004) model of European human population size history, a selective sweep causes a very similar reduction in diversity and skew away from intermediate allele frequencies (mean *π*: 6.49, and mean *D*: -2.27, versus neutral expectations of 47.25 and -0.67, respectively). With increasing genetic distance from a sweep under the bottleneck model, *π* and Tajima’s *D* recover in a similar manner as under constant population size.

Exponential growth is often used to model recent population expansions in lieu of instantaneous population size change, and indeed such growth appears to be a key feature of human population history (Fagundes *et al.* 2007; Gravel *et al.* 2011; Tennessen *et al.* 2012). We therefore examined a model based on the same parameterization of strong growth from Gravel et al.’s (2011) estimated European model but omitting the population contractions and most ancient expansion (Figure 1C). Under this model we find that the impact of positive selection is less severe than under either the equilibrium or bottleneck models: *π* is reduced only 4.5-fold from a neutral expectation of 6.57 to 1.45 when a sweep occurs immediately adjacent to the sampled locus, while Tajima’s *D* only drops from -2.06 to -2.57. This demonstrates in part how strong population growth in the absence of selection creates patterns of variation that are very similar to selective sweeps, so much so that sweeps might be more difficult to detect (Schrider and Kern 2016). Finally, we examined a three-epoch model with a population contraction and then subsequent exponential growth. Currently, this contraction-then-growth model is used to represent non-African human population size histories (e.g. Fagundes *et al.* 2007; Gutenkunst *et al.* 2009; Gravel *et al.* 2011; Tennessen *et al.* 2012). Our parameterization shown in Figure 1D is a simplified version of the European model from Gravel et al. (2011). Under this contraction-then-growth model, we find that *π* experiences a 7.3-fold reduction (from 40.10 to 5.48 on average) when a recent sweep is adjacent to the sampled locus—more similar in scale to the reduction in diversity under the constant-size and bottleneck models than the growth model. Tajima’s *D* also experiences a dramatic reduction when *c*/*s*=0 (from a mean of 0.22 to -2.32). As *c*/*s* increases, both *π* and *D* gradually recover toward their neutral expectations as in the other scenarios.

### Demographic parameter estimates are biased by positive selection

We sought to quantify the impact of positive selection on demographic parameter estimation under our bottleneck, growth, and contraction-then-growth models. First, we simulated population samples experiencing no selection and asked how well we could recover the true parameters of the model using diffusion approximations to the SFS via the ∂a∂i software package (Gutenkunst *et al.* 2009), or with a set of commonly used summary statistics via ABC (Thornton 2009; Csilléry *et al.* 2012); we address the effects of selection on PSMC later, as this method requires a different sampling scheme. Briefly, we used both of these methods to fit the focal demographic model to data sampled from 500 unlinked simulated loci, and repeated this process on 100 replicate simulated “genomes” (Methods). We then gradually increased the value of *f*, the fraction of these sampled loci linked to hard selective sweeps (within a distance of *c*/*s* ≤ 1.0; Methods). At values of *f* ranging from 0 to 1, we repeated parameter estimation to assess the extent to which a given amount of selection biases our inference.

### Population bottleneck

When using ∂a∂i to infer the optimal set of parameters of a bottlenecked population (Figure 1B) experiencing no positive selection, our estimates were quite accurate (Figure 2): our average parameter estimate for the ancestral effective population size, *Ne_A_*, was 10,044 individuals (a 0.4% deviation from the true parameter value); our mean estimate for the time of recovery from the bottleneck, *T_R_*, was 3,126 generations ago (4.2% deviation); our estimated effective population size during the bottleneck, *Ne_B_* was 1,988 individuals (0.6% deviation) on average; and our mean estimate of the present-day effective population size, *Ne*_0_, was 20,476 (2.4% deviation). Moreover, our inferences were fairly consistent, with most parameter estimates being fairly close to the true value (Figure 2). However, while repeating this analysis with increasing numbers of loci linked to a selective sweep, our parameter estimates became increasingly biased. Even a small value of *f* produces significant underestimates of the population sizes *Ne*_0_ and *Ne_B_* (Figure 2). For example, the mean inferred value of Ne_*B*_ decreases to 1,750 when *f*=0.2 (an 12.5% underestimate), to 1,410 when *f*=0.5 (29.5% underestimate), and to 671 when *f*=1.0 (66.4% underestimate). A more subtle but consistent downward bias of *Ne*_0_ also appears with increasing *f*: *Ne*_0_ is estimated at 19,662 at *f*=0.2 (1.7% underestimate), 18,364 when *f*=0.5 (8.2% underestimate), and 15,427 when *f*=1.0 (22.9% underestimate). By contrast, estimates of the ancestral effective population size (*Ne_A_*) and the time since the recovery (*T_R_*) are largely unaffected unless *f* is fairly high (≥0.8), in which case the values of these two parameters are somewhat overestimated (Figure 2).

**Fig. 2.**
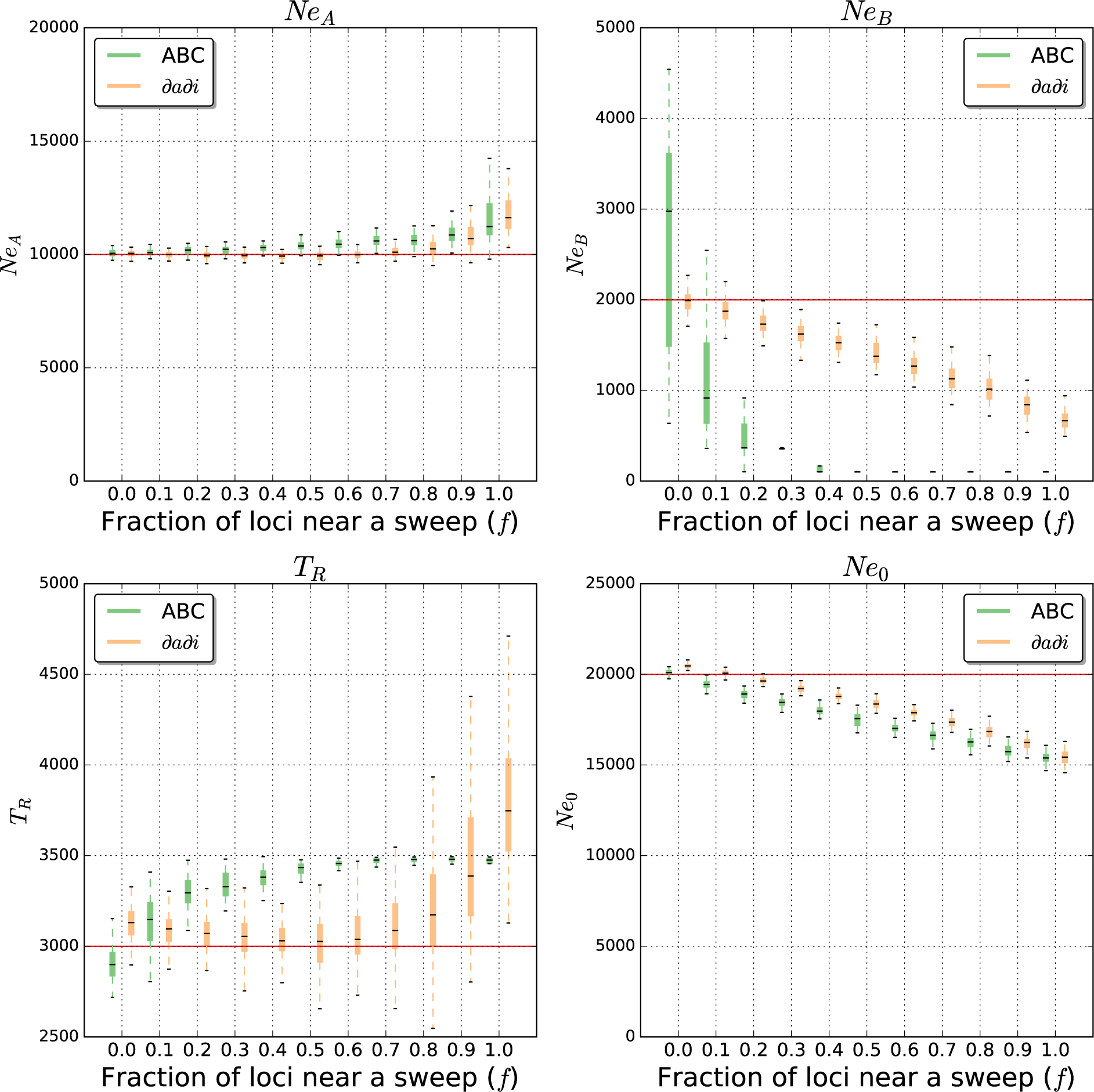
Bottleneck model parameter estimates from ∂a∂i and ABC. Parameter estimation was performed on simulated data sets either evolving neutrally, or with some fraction of loci (*f*) used for inference linked to a selective sweep. Each box plot summarizes estimates from 100 replicates for each scenario. Note that *T_B_*, the bottleneck onset time, is absent from this figure because it was fixed to the true value (Methods).

Like inference using the SFS (i.e. ∂a∂i), our ABC procedure was able to infer the true parameters with minimal bias when run on simulated population samples experiencing no positive selection: the mean estimates were 10,062 for *Ne_A_* (0.6% difference from true value), 2,910 for *T_R_* (3.0% difference), 2,633 for *Ne_B_* (31.7% difference), and 20,097 for *Ne*_0_ (0.5% difference). However, we note that the *Ne_B_* estimate was fairly inconsistent (with the middle 50% of estimates ranging from 1,482 to 3,596), while other parameter estimates exhibited much lower variance. When positive selection is introduced, we obtain significantly biased estimates of all parameters when *f*≥0.2 except for *Ne_A_*. These biases are in the same direction as observed using ∂a∂i (underestimates for *Ne*_0_ and *Ne_B_*, and overestimates for *Ne_A_* and *T_R_*), but typically larger in magnitude. Indeed, for *f*≥0.4, our estimates of *Ne_B_* and *T_R_* approach the boundaries of our prior parameter ranges (the upper bound of 3,500 for *T_R_* and the lower bound of 100 for *Ne_B_*, respectively); in these cases we are inferring a very short but extreme bottleneck. Thus, for our bottleneck model, ABC based on our set of summary statistics appears to be more sensitive to selection than ∂a∂i. Overall, the presence of positive selection seems to cause both methods to overestimate the extent of population contraction, and to underestimate the degree of recovery from the bottleneck. For the simulated datasets used in these analyses, whenever a locus was linked to a selective sweep, the distance from the sweep, *c*/*s* was drawn uniformly between 0 and 1. We also repeated these analyses when fixing the value of *c*/*s*, and in Figure S1 we show our distribution of parameter estimates obtained using both ∂a∂i and ABC on 111 different combinations of *f* and *c*/*s*. This figure demonstrates that, for a given fraction of neutral loci linked to a selective sweep, decreasing the genetic distance to the sweep increases bias, as expected.

Note that for our ABC inference we examined only the means of several population genetic summary statistics (Methods). Including the variances caused estimates to behave non-monotonically, because whenever *f* is not equal to one or zero the distribution of summary statistic values is a mixture of two models, and therefore has inflated variance, resulting in less accurate parameter estimation. We also show our parameter estimates when including variances in Figure S1. In Figure S2, we examine the effect of a recurrent hitchhiking scenario (Methods), finding that selection generally downwardly biases our estimates of *Ne_A_*, *Ne_B_*, and *Ne*_0_ (though for ABC the relationship between *f* and the inferred value is non-monotonic). For *T_R_*, selection decreases estimates from ∂a∂i and ABC using statistic means and variances, but tends to inflate estimates for ABC using only means.

### Population growth

Next, we examined the impact of positive selection on parameter estimates for our model of population growth (Figure 1C). When our simulated genomes experienced no recent selective sweeps, we again achieved good accuracy when using ∂a∂i (Figure 3): our mean estimates of *Ne_A_, T_G_*, and *Ne*_0_ were 1,042 (0.9% difference from true value), 952 (3.5% difference), and 36,819 (2.6% difference), respectively. Increasing *f* again biases our estimates, but the effect is subtler than for the bottleneck case. This is probably a consequence of the reduced scale of the impact of positive selection on flanking variation under this model relative to the bottleneck model (Figure 1). The most notable pattern that we observe for this model is that *T_G_* decreases with increasing *f*, while the population size estimates are largely unaffected: when *f*=0.5 our average estimate is 815 (11.4% difference from true value), versus 735 when *f*=0.8 (20.2% difference), and 679 when *f*=1.0 (26.2% difference). In other words, widespread selective sweeps will cause one to infer slightly more recent but more pronounced exponential growth. When *c*/*s* is relatively small, our error rates are substantially higher (Figure S3). Thus, stronger positive selection could still seriously impair ∂a∂i’s demographic inferences under this population size history.

**Fig. 3.**
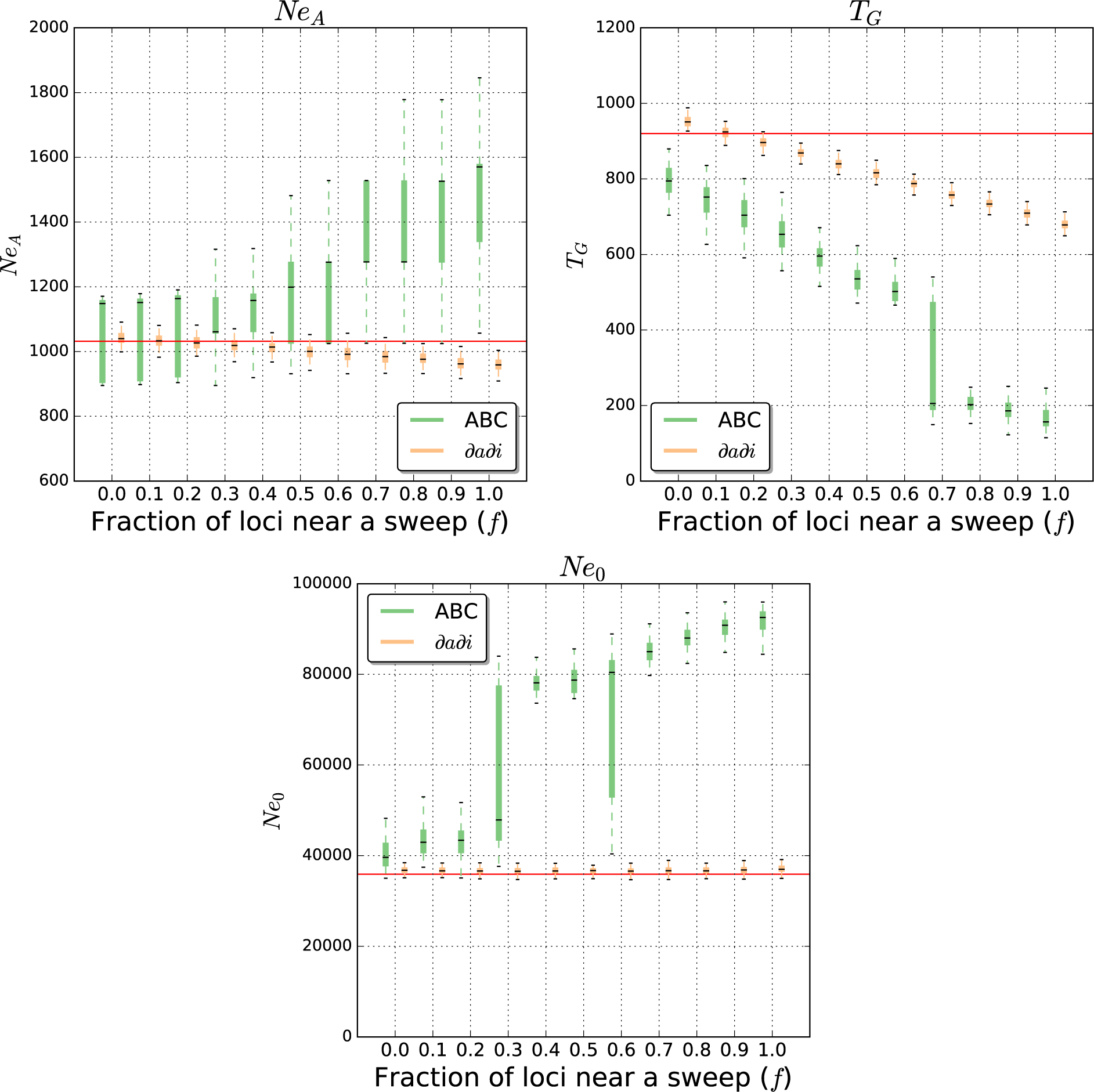
Exponential growth model parameter estimates from ∂a∂i and ABC. Parameter estimation was performed on simulated data sets either evolving neutrally, or with some fraction (*f*) of loci used for inference linked to a selective sweep. Each box plot summarizes estimates from 100 replicates for each scenario.

We then used ABC to perform parameter estimation under the growth model. In the neutral case, our estimated parameters were largely concordant with the true values, with the exception of some bias observed for *T_G_* (mean estimate of 793, which is 13.8% below the true value). Our estimates of *Ne_A_* were also far more dispersed than those obtained from ∂a∂i (Figure 3). Further, unlike our estimates with ∂a∂i, increasing the value of *f* substantially biases our ABC estimates. For example, *Ne*_0_ is 40,215 when *f*=0 (12.0% greater than the true value), but increases to 73,315 when *f*=0.5 (an overestimate of 104.2%), 87,702 when *f*=0.8 (an overestimate of 144.3%), and 91,725 when *f*=1.0 (an overestimate of 155.5% plus a dramatic increase in variance). The degree to which *T_G_* is underestimated also increases with *f*: the average estimate is 705 at *f*=0.2 (23.4% below the true value), 519 at *f*=0.5 (43.5% bias), 233 at *f*=0.8 (74.7% bias), and 167 at *f*=1.0 (81.2% bias). Again, we demonstrate the effect of varying the distance *c*/*s* of sampled loci from the selective sweep, as well as the effect of performing ABC on the variances of summary statistics in addition to their means, in Figure S3. Overall, we observe that under our growth model positive selection will cause inferences of more recent, faster population growth, with this effect being far subtler when using ∂a∂i than ABC with our set of summary statistics. When recurrent hitchhiking model (Methods), we find that selection downwardly biases our estimates of *Ne_A_* from each method, and dramatically inflates estimates of *Ne*_0_ for ABC using means and variances (Figure S4).

### Population contraction followed by growth

Finally, we assessed our ability to recover the parameters of our contraction-then-growth model (Figure 1D) with increasing amounts of positive selection. Without selection, ∂a∂i estimates *Ne_A_, T_G_*, and *Ne*_0_ with reasonably high accuracy (Figure 4): 14,767 on average for *Ne_A_* (2.0% over the true value), 864 for *T_G_* (6.1% under the true value), and 37,922 for *Ne*_0_ (5.6% over the true value). However, *T_C_* and *Ne_C_* are substantially overestimated at 2,511 (23.1% over the true value) and 1,339 on average (29.7% over the true value), respectively. As we increase *f*, our estimates of *Ne_A_, T_C_*, and *Ne_C_* are inflated, *T_G_* is increasingly underestimated, and *Ne*_0_ is largely unaffected. The effect on *T_C_* is the largest, resulting in a massive increase with *f*: our estimate is 3,719 when *f*=0.2 (an overestimate of 82.3%), 10,467 when *f*=0.5 (overestimate of 413.1%), and 13,728 when *f*=0.8 (overestimate of 572.9%). *Ne_A_* and *Ne_C_* estimates also increase dramatically with *f*: to 49,431 (an overestimate of 241.5%) and 2,415 (an overestimate of 134.0%) when *f*=0.5, respectively, while *T_G_* on the other hand decreases to 580 when *f*=0.5 (underestimate of 36.9%). Thus, positive selection typically results in our ∂a∂i-estimated demographic model to have more protracted population contraction, with larger initial and contracted population sizes.

**Fig. 4.**
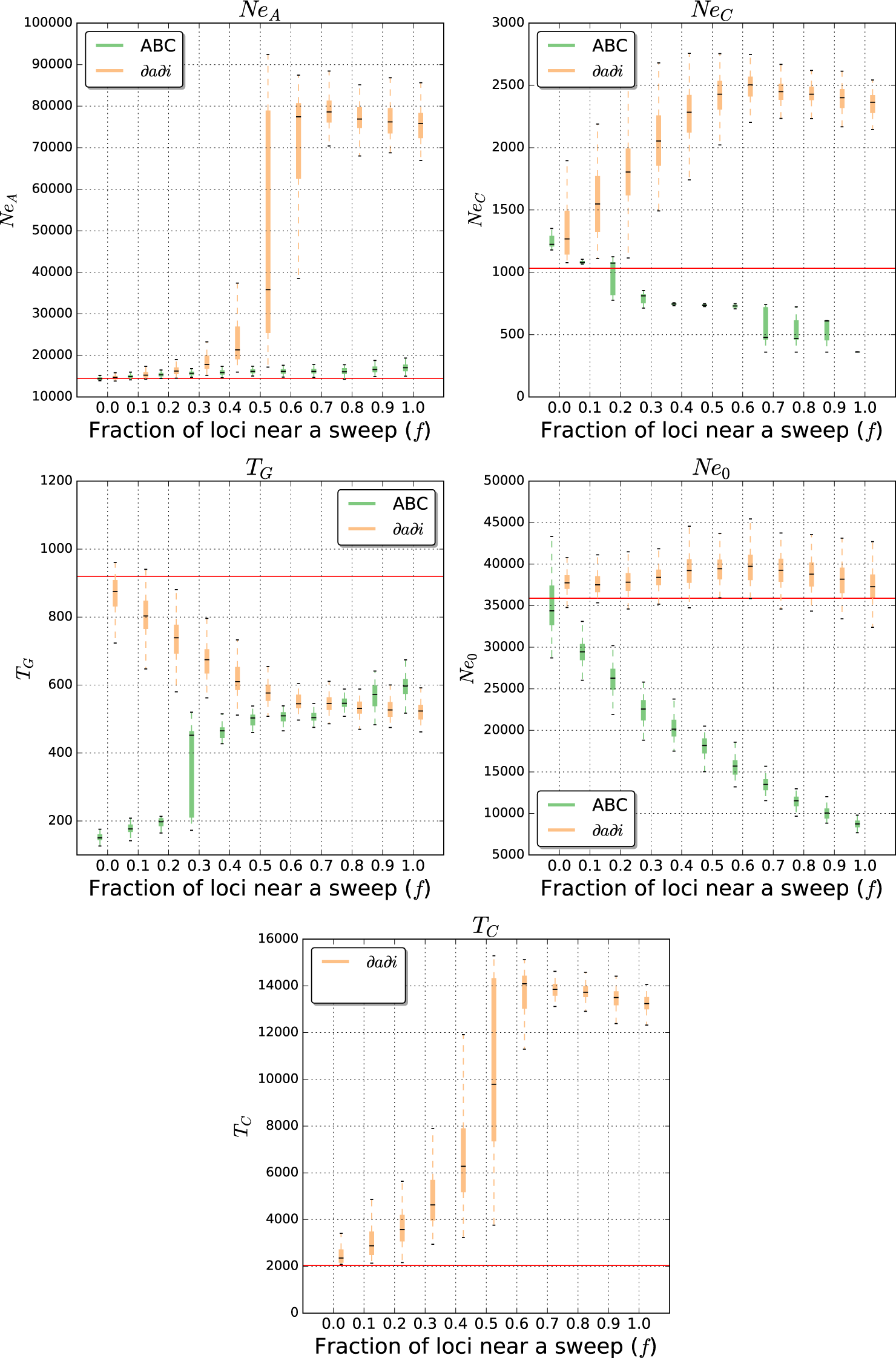
Contraction-then-growth model parameter estimates from ∂a∂i and ABC. Parameter estimation was performed on simulated data sets either evolving neutrally, or with some fraction of loci (*f*) used for inference linked to a selective sweep. Each box plot summarizes estimates from 100 replicates for each scenario. Note that when performing ABC, time of population contraction (*T_C_*), was fixed to the true value and therefore this parameter is only shown for ∂a∂i.

When repeating these analyses using ABC given our set of summary statistics (Figure 4), we find that under neutrality *T_G_* is grossly underestimated, *Ne_C_* is slightly overestimated, and *Ne_A_* and *Ne*_0_ are estimated with greater accuracy (14,415 or 0.4% under the true value, and 35,017, or 2.5% under the true value, respectively). Thus, we infer a more protracted but slightly less severe population contraction than the true population size history. Even so we proceed to characterize what the effect of linked selection on parameter estimates using ABC as before. Indeed our estimates become more biased as we add increasing amounts of positive selection. Most notably, *Ne*_0_ exhibits a substantial downward bias as we increase *f*, and is estimated at 26,178 when *f*=0.2 (27.1% underestimate), 18,101 when *f*=0.5 (49.6% underestimate), and 8,794 when *f*=1.0 (75.5% underestimate). Also, as *f* becomes large *Ne_C_* shifts from being slightly overestimated to significantly underestimated, and estimates of *Ne_A_* become slightly upwardly biased. Thus we find that under this model positive selection again biases parameter estimates, though not in the same manner for ∂a∂i and ABC: while ∂a∂i infers a longer phase of reduced population size along with an inflated ancestral size and less severe contraction, our ABC procedure infers a more severe contraction followed by a less complete recovery. We show our inference results on the full grid of *c*/*s* and *f* values, as well as when including variances of summary statistics, in Figure S5. In Figure S6, we show estimates under recurrent hitchhiking, where we observe that ∂a∂i’s estimates of *T_C_* are inflated, while *Ne_A_* is also skewed, with ∂a∂i producing underestimates with lower values of *f* and overestimates when *f*≥0.8. ABC under recurrent hitchhiking produces underestimates of *Ne_A_*, and oscillates between downwardly and upwardly biased inferences for *Ne*_0_. In general, demographic parameter estimation under our contraction-with-growth scenario in the presence of recurrent hitchhiking leads to very poor estimates under a wide range of *f*.

### Effect of positive selection on population size history inference using PSMC

The pairwise sequentially Markovian coalescent (PSMC) is a widely used method that infers a discretized history of population size changes from a single recombining diploid genome (Li and Durbin 2011). Such inference is possible because coalescence times between the two allelic copies in a diploid, which are governed by the effective population size, will change at the breakpoints of historical recombination events, and the resulting distribution of coalescence times across the genome thus contains information about population size history. However this method necessitates sampling a large stretch of a recombining chromosome. In order to test the impact of positive selection on inferences from PSMC, we simulated constant-size populations of 10,000 individuals, sampling a 15 Mb chromosomal region from two haploid individuals. We performed 100 replicates of this simulation for each of four scenarios (Methods): the standard neutral model; a population experiencing one fairly recent sweep (reaching fixation 0.2×*Ne* generations ago) somewhere in this region; a population experiencing three recurrent sweeps (fixed 0, 0.2, and 0.4×*Ne* generations ago); and a population experiencing five sweeps (0, 0.1, 0.2, 0.3, and 0.4×*Ne* generations ago). We find that under neutrality, very little population size change is inferred on average (though there is a fair bit of variance; Figure 5A). However, when there has been only a single selective sweep, a population bottleneck near the time of the sweep is inferred, in which the population contracts to approximately one-half of its original size before recovering to greater than twice its original size (Figure 5B). When there have been three or five recurrent selective sweeps the inferred population contraction becomes increasingly severe (Figure 5C-D). In the five-sweep case, we typically infer a contraction down to roughly one-fourth of the original size, though the subsequent growth phase is less dramatic than that inferred from the one- and three-sweep cases. We also find that a recurrent hitchhiking regime substantially biases the inferred population size history when the rate of hitchhiking becomes appreciable (i.e. ≥1 sweep per 15 Mb region every 2*N* generations; see Figure S7). In particular, recurrent hitchhiking leads to a case where the population is in constant recovery from a loss of polymorphism, thus even though population size has remained constant, PSMC infers a population size history that looks like population growth. Selection also biases PSMC’s inference when the population size varies over time: in Figure S8 we show that under our contraction-then-growth model, the duration of the bottleneck is greatly overestimated as sweeps are added to the scenario, with a more ancient contraction inferred. In addition, panels C and D of Figure S8 suggest that the extent of exponential growth during the recovery phase is underestimated when very recent sweeps have occurred. Thus, we find that positive selection can dramatically skew population size histories deduced by PSMC.

**Fig. 5.**
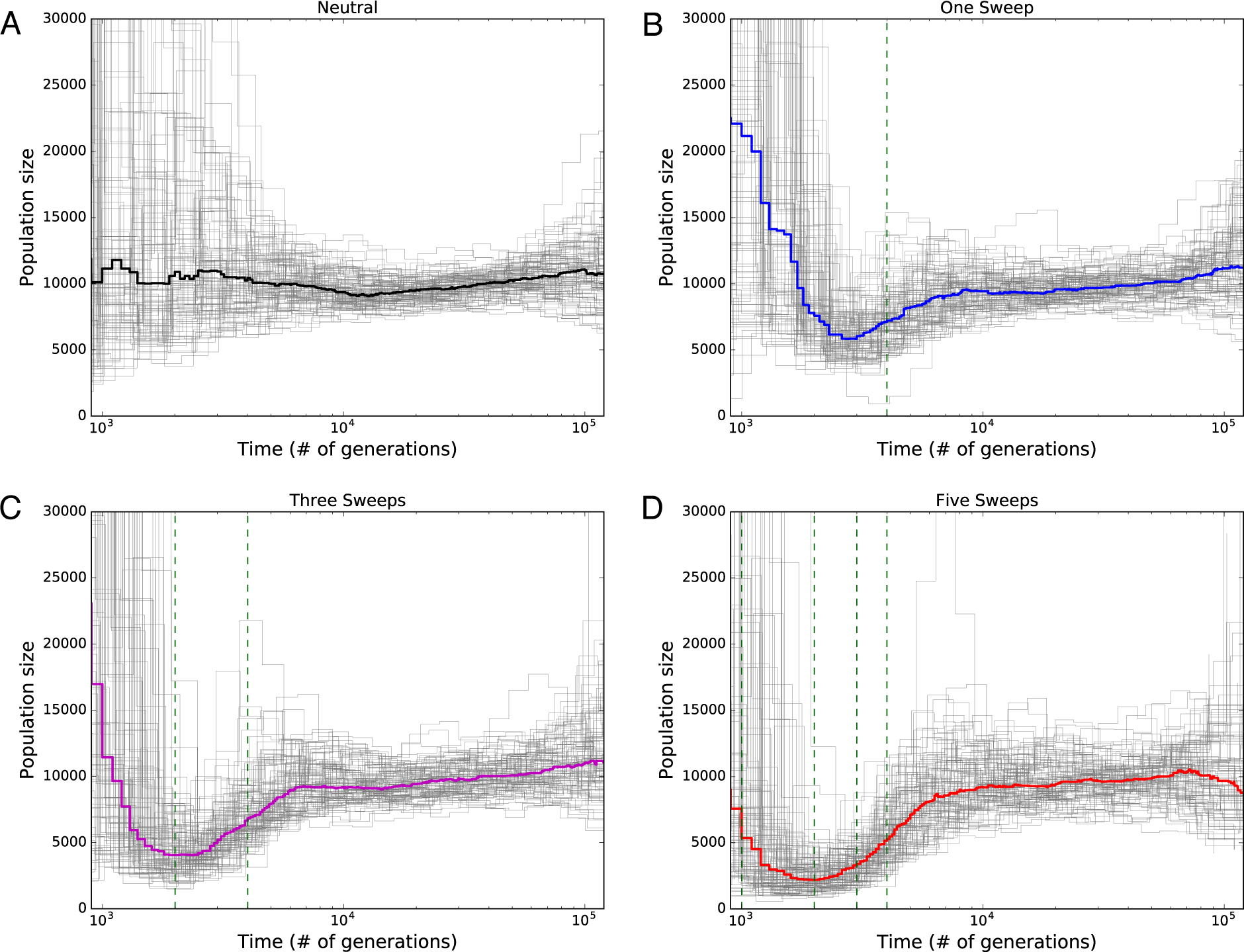
Population size histories inferred by PSMC. Inferred size histories for each of 100 replicate simulations are shown as thin gray lines, and the median across all replicates is shown as the thicker line. In all cases the simulated population’s size was constant throughout. (*A*) Population size histories inferred from neutral simulations. (*B*) Inferences from simulations with one selective sweep, for which the fixation time is shown as a dashed green vertical line. (*C*) Inferences from simulations with three recurrent selective sweeps. Fixation times for the two older sweeps are shown as dashed green vertical lines, while the most recent sweep fixed immediately prior to sampling. (*D*) Five recurrent selective sweeps, with fixation times for the four oldest shown as dashed vertical lines; again, the most recent sweep fixed immediately prior to sampling.

### Positive selection produces spurious support for non-equilibrium demographic histories

Demographic inference methods are often used not only to infer parameters of a model, but increasingly to select the best fitting among several competing models (e.g. Adams and Hudson 2004; Fagundes *et al.* 2007; Duchen *et al.* 2013). To ask whether positive selection might affect the outcome of demographic model selection, we simulated genomes with constant population size, again sampling loci for which some fraction, *f*, is located within *c*/*s* ≤ 1 of a selective sweep. We then performed model selection among our four demographic histories (Figure 1) using both ∂a∂i and ABC (Methods).

Prior to performing model selection with ∂a∂i, we first examined the degree of support for each model when fit to each dataset using the Akaike information criterion (AIC). Examining the differences in AIC between models, we found that even a moderate number of selective sweeps will cause non-equilibrium demographic scenarios to have far stronger support than the true equilibrium history (Figure S9). This is especially so for the bottleneck and contraction-then-growth models, which achieve better support than the equilibrium model even at small values of *f*. For example, when *f*=0.2 the bottleneck model receives an AIC lower than the equilibrium model in 90% of cases, and the contraction-then-growth model has a lower AIC 72% of the time (Figure S9). By contrast, the pure growth model is supported to a lesser extent (a lower AIC in 54% of cases), and occasionally failed to optimize properly, settling on a very low-likelihood parameterization—an indication of a poorly fitting model. The better fit of the bottleneck and contraction-then-growth models is likely because they better model the genealogy of a region experiencing a selective sweep: much of the ancestral variation flanking the selected site is removed during the sweep (analogous to contraction), while being replaced by the subset of alleles within the rapidly expanding class of individuals containing the selected mutation (analogous to expansion).

We conducted formal model selection as described in the Methods, conservatively selecting the equilibrium model unless one of the other models had an AIC at least 50 units lower. We note that it would be preferable to perform parametric bootstraps from competing models to compare the distributions of AIC values, but in the interest of computational efficiency we instead choose this heuristic. Even with this conservative cutoff, we selected a non-equilibrium model for 15% of simulated data sets with *f*=0.2, for 47% of datasets with *f*=0.3, and for 91% of datasets when *f*=0.6 (Table 1). Thus even if a minority of loci are linked to a recent selective sweep then SFS-likelihood based approaches may prefer the wrong demographic model. Interestingly, in every case where a non-equilibrium model was the unambiguous best fit to the data this model choice was the bottleneck scenario. We also performed the same model selection procedure on constant-size populations experiencing recurrent hitchhiking (Methods), finding that the neutral model was rejected in all cases where some fraction of the genome experienced selection (*f* ≥0.1; Table S1).

**Table 1:** The fraction of simulated data sets for which each demographic model was selected by ∂a∂i.

| $f$ | Bottleneck | Growth | Contraction-then-growth | Non-equilibrium (ambiguous) | Equilibrium |
| --- | --- | --- | --- | --- | --- |
| 0 | 0.0% | 0.0% | 0.0% | 0.0% | 100.0% |
| 0.1 | 1.0% | 0.0% | 0.0% | 1.0% | 98.0% |
| 0.2 | 5.0% | 0.0% | 0.0% | 10.0% | 85.0% |
| 0.3 | 14.0% | 0.0% | 0.0% | 33.0% | 53.0% |
| 0.4 | 41.0% | 0.0% | 0.0% | 55.0% | 4.0% |
| 0.5 | 77.0% | 0.0% | 0.0% | 23.0% | 0.0% |
| 0.6 | 91.0% | 0.0% | 0.0% | 9.0% | 0.0% |
| 0.7 | 100.0% | 0.0% | 0.0% | 0.0% | 0.0% |
| 0.8 | 99.0% | 0.0% | 0.0% | 1.0% | 0.0% |
| 0.9 | 100.0% | 0.0% | 0.0% | 0.0% | 0.0% |
| 1 | 100.0% | 0.0% | 0.0% | 0.0% | 0.0% |

Next, we performed model selection on our constant-size population samples using ABC (Methods). For each of these datasets, we estimated Bayes factors for each pairwise comparison of demographic models. Again, we find that non-equilibrium demographic models may begin to receive stronger support than the constant-size model once a sizable fraction of loci are linked to selective sweeps. For example, when *f*=0.4, the bottleneck model has nominally stronger support (Bayes factor >1) than the equilibrium model for 55% of datasets, the growth model has stronger support than equilibrium in 9% of datasets, and the contraction-then-growth model has stronger support in 4% of datasets (Figure S10). When *f* is increased to 0.8, we observe even stronger support for non-equilibrium models, with 100% of the bottlenecks datasets, 79% of the contraction-the-growth datasets, and 26% of the growth model datasets having a Bayes factor > 1 when compared to the constant-size model. We used these Bayes factors to perform model selection in a manner similar to our analysis with ∂a∂i, conservatively selecting the equilibrium model if there was no alternative model that was a significantly better fit to the data (i.e. having a Bayes factor relative to the equilibrium model of ≥20). Again, we find that even if a minority of loci are linked to a sweep, then there is a substantial probability that the constant-size model will not be selected: for 6% of datasets we select a non-equilibrium model when *f*=0.2, for 34% of datasets with *f*=0.3, and for 99% of datasets when *f*=0.6 (Table 2). As with ∂a∂i-optimized models, we found that in every instance where we were able to unambiguously select a single non-equilibrium demographic history as the best fit we chose the bottleneck model. When we include the variances of our set of summary statistics in our ABC procedure, we find that non-equilibrium models are strongly supported in an even higher proportion of simulated data sets, though in this case we typically select the contraction-then-growth model rather than the bottleneck model (Table S2). Similarly, when performing our ABC model selection on datasets with recurrent hitchhiking (Methods), we find that the constant-size model was nearly always rejected in the presence of recurrent sweeps (see Table S3 for model selection using means of summary statistics and Table S4 for means and variances).

**Table 2:** The fraction of simulated data sets for which each demographic model was selected by ABC.

| $f$ | Bottleneck | Growth | Contraction-then-growth | Non-equilibrium (ambiguous) | Equilibrium |
| --- | --- | --- | --- | --- | --- |
| 0 | 0.0% | 0.0% | 0.0% | 0.0% | 100.0% |
| 0.1 | 0.0% | 0.0% | 0.0% | 1.0% | 99.0% |
| 0.2 | 0.0% | 0.0% | 0.0% | 6.0% | 94.0% |
| 0.3 | 2.0% | 0.0% | 0.0% | 32.0% | 66.0% |
| 0.4 | 25.0% | 0.0% | 0.0% | 45.0% | 30.0% |
| 0.5 | 88.0% | 0.0% | 0.0% | 2.0% | 10.0% |
| 0.6 | 99.0% | 0.0% | 0.0% | 0.0% | 1.0% |
| 0.7 | 100.0% | 0.0% | 0.0% | 0.0% | 0.0% |
| 0.8 | 100.0% | 0.0% | 0.0% | 0.0% | 0.0% |
| 0.9 | 100.0% | 0.0% | 0.0% | 0.0% | 0.0% |
| 1 | 100.0% | 0.0% | 0.0% | 0.0% | 0.0% |

## DISCUSSION

It is well known that natural selection profoundly affects genealogies and therefore patterns of genetic polymorphism (Kaplan *et al.* 1989; Hudson and Kaplan 1994), thus it is reasonable to expect that linked selection will bias demographic inference that assumes strict neutrality of population genomic data. Indeed, background selection has recently been shown to skew demographic inferences using the site frequency spectrum (Ewing and Jensen 2016). Here, we show through extensive simulation that positive selection can severely impair demographic model selection and parameter estimation based on the SFS, summary statistics of variation, and reduced approximations of the ancestral recombination graph (i.e. PSMC). The extent to which this is so depends on the fraction of genetic loci examined during inference that are affected by a recent sweep, and the ratio of the genetic distance between the locus and the target of selection to the selection coefficient (*c*/*s*).

When the fraction of loci affected by linked selection is low, we have shown that point estimates of population parameters estimated under the correct demographic scenario are reasonably accurate using both SFS-based inference and ABC with summary statistics (figs. 2-4)—the exact fraction, however, depends on the model in question. However, unless *f* is quite low our results indicate that when model selection is applied using either SFS-based or ABC inference, linked selection can bias model choice (Tables 1 and 2). In many of our simulated datasets we have assumed that loci linked to sweeps are on average a *c*/*s* distance of 0.5 away from the sweep (i.e. drawn uniformly from between 0 and 1). In real genomes this may correspond to quite a large physical distance: for instance if we assume a selection coefficient of 0.05 (i.e. selection as strong as in our simulations) and a crossover rate of 2 cM/Mb (similar to estimates in Drosophila; e.g. Comeron *et al.* 2012) this corresponds to a physical distance of 1.25 Mb. If instead we assume a crossover rate of 1 cM/Mb (similar to estimates from humans; e.g. Kong *et al.* 2010), this corresponds to a physical distance of 2.5 Mb. While we have assumed a fairly high value of *s* that may not be representative of all selective sweeps (e.g. Li and Stephan 2006; Jensen *et al.* 2008), known sweeps in humans may often have selection coefficients fairly close to 0.05 (Peter *et al.* 2012).

Thus, even if there are a small number of recent selective sweeps, the majority of the genome may nonetheless be sufficiently impacted by linked selection to produce biased demographic inferences. For example, if a human population experienced 1,200 recent sweeps fairly evenly spaced across the genome (the equivalent of one recent sweep in ∼5% of genes), every site in the genome would be within a *c*/*s* distance of 0.5 (assuming *s*=0.05) from the nearest sweep (i.e. *f* = 1.0). This is likely an unrealistic scenario in human populations, where positive selection is perhaps less common (Hernandez *et al.* 2011). However, some have argued that selection may be pervasive in the human genome as well (Boyko *et al.* 2008; Enard *et al.* 2014), and certainly humans show many adaptations to local environments (e.g. Li *et al.* 2007; Perry *et al.* 2007; Tishkoff *et al.* 2007; Barreiro *et al.* 2008; Bryk *et al.* 2008). Thus while it seems unlikely that every site in the genome would be affected by linked selection, some currently unknown subset of sites must be and thus we should expect demographic estimation to be subject to some degree of bias (perhaps negligible in well-designed studies; see below). In Drosophila, each genomic position would be closely linked to a sweep (*c*/*s* distance of <0.5) if there were as few as 120 evenly spaced recent sweeps (again assuming *s*=0.05). This is a very small number indeed, equivalent to a single recent sweep affecting <1% of all genes. Indeed in Drosophila the fraction of loci affected by recent positive selection may be quite large (Begun *et al.* 2007; Langley *et al.* 2012). Numerous studies have estimated that the fraction of adaptive amino acid substitutions in *D. melanogaster* is considerable, with estimates ranging from 10-50% (Smith and Eyre-Walker 2002; Bierne and Eyre-Walker 2004; Langley *et al.* 2012; Mackay *et al.* 2012). Direct estimates of the rate of recurrent hitchhiking in Drosophila populations also indicate a substantial flux of adaptive substitution, with the population effective sweep rate varying between 1.1×10^-5^ to 2.6×10^-3^ sweeps per base pair per 2*N* generations (Li and Stephan 2006; Andolfatto 2007; Macpherson *et al.* 2007; Jensen *et al.* 2008). For instance, Li and Stephan (Li and Stephan 2006) in analyzing African population data estimated that approximately 160 strong selective sweeps had occurred in the recent history on the X chromosome alone, an estimate that was in close agreement to that from Jensen et al. (2008) using different methods. Our simulations of recurrent hitchhiking with rates of sweeps roughly in line with those of Li and Stephan (2006) and Jensen et al. (2008), led to biased demographic inference in most cases. Positive selection may therefore be particularly troublesome for demographic inference in Drosophila and other organisms were adaptive natural selection is similarly pervasive. In sum then, the 10-fold larger number of sweeps (under our parameterization) required to produce the same level of bias suggests that the confounding effect of demography in humans may be less problematic than in Drosophila. However, given uncertainty in the number, location, strength, and type (see below) of selective sweeps we are unable to quantify the extent to which demographic inferences in either species are skewed by adaptation. For instance, if we decrease our assumed value of *s* by an order of magnitude, the number of sweeps required to place each locus in the genome (*f*=1.0) within *c*/*s* of 0.5 of a sweep would have to be increased by an order of magnitude in the examples given above. However we note that in many cases even much lower values of *f* can bias demographic estimates (Figures 2-4). Finally, our goal is not to provide a compelling case for large-scale distortions in current demographic estimates in any single population (e.g. humans or flies) but instead to demonstrate how linked selection can bias inference in an idealized population.

A new and promising class of methods for inferring demographic histories rely on estimating approximations to the ancestral recombination graph (ARG) using a sequentially Markovian coalescent (Li and Durbin 2011; Schiffels and Durbin 2014). In particular, because PSMC is applied to a single diploid genome, it has been used to infer population size histories in numerous species for which one or more genome sequences are available, in many cases finding support for large changes in population size (Groenen *et al.* 2012; Albert *et al.* 2013; Prado-Martinez *et al.* 2013; Zhan *et al.* 2013; Freedman *et al.* 2014; Green *et al.* 2014; Wallberg *et al.* 2014;Auton *et al.* 2015; Lamichhaney *et al.* 2015). Our findings suggest that natural selection may alter the shape of, and inflate the degree of change in, these inferred histories. Indeed because of the specific way in which a sweep perturbs the ARG locally during the coalescent history of a chromosome, PSMC inference on regions that have experienced one or a few sweeps in the past may lead to erroneous estimation of a population bottleneck. If sweeps continue until the present day, PSMC inference might appear to support a population contraction rather than a bottleneck, though this may very well be a result of PSMC having lower power for very recent population dynamics (Li and Durbin 2011). Under a truly recurrent sweep model (Stephan *et al.* 1992) PSMC infers populations size history that resemble growth, as the genomic region is in a case of steady recovery from the last sweep to have occurred (see Figure S7).

Our results are broadly concordant with a recent examination of the ability of the McDonald–Kreitman (or MK) test (McDonald and Kreitman 1991) to infer the fraction of substitutions that were adaptive (*α*) under a simulated recurrent hitchhiking scenario with constant population size (Messer and Petrov 2013). Their study found that Eyre-Walker and Keightley’s DFE-alpha method (Eyre-Walker and Keightley 2009), which simultaneously estimates *α*, the distribution of fitness effects, and a two-epoch population size history, incorrectly inferred the presence of population size changes (Messer and Petrov 2013). It is therefore reasonable to assume that positive selection could have a substantial confounding effect on a variety of population genomic methods for demographic inference in practice, beyond those considered here. In the empirical literature, numerous recent studies of demographic history have found support for contractions and recent expansions of natural populations (Thornton and Andolfatto 2006; Fagundes *et al.* 2007; Gravel *et al.* 2011; Tennessen *et al.* 2012; Duchen *et al.* 2013). While such population size changes are probably common, and our results do not call the major findings of these studies into question, they do suggest that natural selection exaggerates the inferred intensity of these changes.

Our study has examined only a single model of adaptive natural selection, and therefore has several limitations. Throughout we have assumed that positive selection occurs only through completed hard selective sweeps. Indeed soft sweeps (Innan and Kim 2004; Hermisson and Pennings 2005; Pennings and Hermisson 2006; Garud *et al.* 2015) and partial sweeps (Hudson *et al.* 1994; Sabeti *et al.* 2002; Voight *et al.* 2006), may be widespread, and differ in their effects on linked polymorphism (Orr and Betancourt 2001; Meiklejohn *et al.* 2004; Przeworski *et al.* 2005; Teshima *et al.* 2006; Schrider *et al.* 2015; Vy and Kim 2015). Polygenic selection, in which alleles at several different loci underlying a trait under selection will experience a change in frequency, is also thought to be widespread (Pritchard *et al.* 2010; Berg and Coop 2014). Such polygenic adaptation is known to leave its own unique signature on patterns of population genetic variation (Berg and Coop 2014). These alternative modes of positive selection could skew demographic inferences in a different manner than what we have observed in this study. Positive selection may also affect estimation of multi-population demographic scenarios: though we did not examine this here, Mathew and Jensen recently showed that selective sweeps will impair parameter estimates for a two-population isolation-with-migration model (Mathew and Jensen 2015). Thus our results, combined with those of Mathew and Jenson (2015), Ewing and Jensen (2016), and Messer and Petrov (2013), strongly suggest that the problem of natural selection skewing demographic inference is a general one.

The observations we have made here also suggest some steps that can be taken to mitigate the impact of positive selection. First, we note that in general ∂a∂i (i.e. SFS-based inference) appears to be somewhat more robust to selection than our ABC approach based on summary statistics. Perhaps this is because ∂a∂i uses an SFS summed across loci, such that more polymorphic regions will have a greater weight on the shape of the SFS (simply because they contribute more observations). Thus the extent to which regions most affected by sweeps contribute to the SFS is diminished implicitly, as these regions will exhibit less variation. Relying on the SFS rather than summaries of variation that to a greater extent depend on the number of segregating sites may therefore reduce selection’s confounding effect on inferred relative population size changes, though estimates of 4*Nµ* and therefore the absolute population size may be biased. We also found that including variances of summary statistics when performing ABC can dramatically inflate error, especially when an intermediate number of loci are linked to sweeps, perhaps because this mixture of two evolutionary models (neutrality and positive selection) inflates the variance. Omitting variances may therefore reduce the confounding effect of selection in some cases.

Finally, we have shown that the proximity of selective sweeps to genomic regions used for inference (as measured by *c*/*s*) has a large effect on the magnitude of bias (supplementary figs. 1-3). Thus, it is of paramount importance to select regions located as far away in genetic distance as possible from genes and other functional DNA elements (Gazave *et al.* 2014)—note that our results imply that short introns and synonymous sites may be poor choices. While this is so, it may not be possible to move far enough away from potential targets of selection to completely eliminate any bias (as discussed above). Moreover, it is essential to omit regions with lowered recombination rates, where the impact of linked selection will be strongest (Begun and Aquadro 1992). As a corollary to this, demographic estimation performed from regions of the genome at differing genetic distances from regions experiencing selection should recover different parameter estimates (Gazave *et al.* 2014). Our results also motivate the challenging task of simultaneous estimation of parameters related to natural selection and demographic history (Eyre-Walker and Keightley 2009; Mathew and Jensen 2015; Sheehan and Song 2016). Until an approach to obtain accurate estimates of demographic parameters in the face of natural selection is devised, population size histories inferred from population genetic datasets from many species may remain substantially biased.

## METHODS

### Simulating demographic and selective histories to test inference methods

To test the robustness of ∂a∂i and ABC to positive selection, we generated coalescent simulations from four different demographic scenarios: 1) a constant population size model; 2) a three-epoch population bottleneck (the European model from Marth *et al.* 2004); 3) a model of recent exponential population growth; 4) and a three-epoch model with a population contraction followed by a period of stasis and then recent exponential population growth (a simplified version of the European model from Gravel *et al.* 2011). We refer to this scenario as the contraction-then-growth model. These models and their parameters are shown in Figure 1.

For each demographic model, we simulated 100 observed genomes experiencing no natural selection, each of which was summarized by a collection of 500 unlinked loci sampled from 200 haploid individuals. We then repeated these simulations while stipulating that a specified fraction of loci (*f*) were linked to a recent selective sweep where the selected mutation reached fixation immediately prior to sampling. The selection coefficient, *s*, for this mutation was always set to 0.05, with a completely additive fitness effect (*h*=0.5). For each simulation including a selective sweep, we specified the genetic distance of the sweep from the sampled locus by the ratio *c*/*s*, where *c* is the crossover rate per base pair multiplied by the physical distance to the sweep, and *s* is again the selection coefficient. We examined values of *f* that were multiples of 0.1 between 0.1 (10% of loci linked to a sweep) and 1.0 (100% of loci linked to a sweep). Values of *c*/*s* examined were multiples of 0.1, raging from 0.0 (the sweep occurred immediately adjacent to the locus being used for inference) to 1.0 (∼4.17 Mb given our value of *s* and our recombination rate; see below). We generated sets of simulations with a given value of *f* by combining the appropriate numbers of neutral simulations and simulated loci linked to a sweep. For each combination of *f* and *c*/*s* (110 combinations in total), we generated 100 sets of 500 unlinked loci.

Next, we repeated these simulations under a recurrent hitchhiking selection scenario where rather than being linked to one recent selective sweep, loci were linked to selective sweeps that arose according to a rate parameter 2*Nλ* sweeps per base pair per 2*N* generations, with the sweep location ranging between 0 and a genetic distance of 4*Ns* away from sequenced locus. We set this rate to 2×10^-5^ per base pair (or a total rate across the entire linked region of map distance 4*Ns* of 100, given that *s*=0.05). While we did not select our parameters to mimic any particular system, we note that our rate and selection coefficient are qualitatively similar to those estimated from Drosophila by Jensen et al. when using a fixed value of *s* rather than a continuous distribution (Jensen *et al.* 2008), though rate estimates vary substantially (see Table 3 from Jensen *et al.* 2008 for examples). For these simulations we did not separately examine multiple values of *c*/*s*. Instead, for a given value of *f*, the appropriate fraction of sampled regions were linked to loci undergoing recurrent sweeps whose location ranged from a *c*/*s* ratio of zero to 2. The remaining fraction, 1-*f*, loci were unlinked from selection.

We also simulated large chromosomal regions to which we applied PSMC. These simulations were of 15 Mb regions with a constant-size population (*Ne*=10,000), from which two individuals were sampled. These 15 Mb regions either experienced no selective sweeps, one selective sweep fixing 0.4×*Ne* generations ago, three selective sweeps (fixing 0.4×*Ne* generations ago, 0.2×*Ne* generations ago, and immediately prior to sampling) or five selective sweeps (0.4×*Ne*, 0.3×*Ne*, 0.2×*Ne*, 0.1×*Ne*, or 0 generations prior to sampling). The location of each sweep was thrown down randomly along the chromosome. For each scenario, 100 replicate simulations were generated. We also simulated 15 Mb regions under our contraction-and-growth scenario with zero sweeps, one sweep (fixing 9000 generations ago, or ∼0.63×*Ne* generations ago), three sweeps (fixing zero, 5000, and 9000 generations ago, or 0×*Ne*, ∼0.35×*Ne*, or ∼0.63×*Ne* generations ago, respectively), or five sweeps (zero, 5000, 7000, and 9000 generations ago, or 0×*Ne*, ∼0.21×*Ne*, ∼0.35×*Ne*, ∼0.49×*Ne*, or ∼0.63×*Ne* generations ago, respectively). Finally, we simulated constant-size populations (*Ne*=10,000) experiencing recurrent hitchhiking at a variety of rates ranging from a rate of 0.01 to 1000 sweeps per 15 Mb per 2*N* generations. Again, all simulated sweeps had *s*=0.05 and *h*=0.5.

For all simulations we used parameters relevant to human populations: a recombination rate of 1.0×10^-8^, (approximately equal to the sex-averaged rate from Kong *et al.* 2010) and a mutation rate of 1.2×10^-8^ (from Kong *et al.* 2012). Simulations were performed with our coalescent simulator discoal (https://github.com/kern-lab/discoal), and example command lines with the appropriate population mutation/recombination rates and population size changes for each demographic scenario are shown in Table S5.

### Parameter estimation and model selection with ∂a∂i

We downloaded version 1.6.3 of ∂a∂i (Gutenkunst *et al.* 2009), which we programmed to optimize the parameters of the bottleneck, growth, and contraction-then-growth models. For each model we used a two-step constrained optimization procedure to find the combination of demographic parameters that have the highest likelihood given the site frequency spectrum measured across all 500 unlinked loci in the simulated genome. First we performed a coarse optimization using the Augmented Lagrangian Particle Swarm Optimizer (Jansen and Perez 2011), and then refined this solution using Sequential Least Squares Quadratic Programming (Kraft 1988). Both of these techniques are implemented in the pyOpt package (version 1.2.0) for optimization in python (Perez *et al.* 2012).

To asses the accuracy of point estimation of parameters in the face of varying amounts of and genetic distances to selective sweeps, we optimized the parameters of each demographic model against each data set simulated under that model, comparing estimated values to the true values. As shown in the Results, this approach was quite successful recovering the true parameter values of each demographic model when applied to data simulated under neutrality. However, one exception was the bottleneck model, for which the optimal solution was typically a shorter but more severe bottleneck than the one we had simulated. We therefore fixed the bottleneck duration to the true value (500 generations), after which ∂a∂i was able to estimate the remaining parameter values with acceptable accuracy.

To assess the support for a given demographic model, we obtained for a simulated data set the likelihood of each demographic model under the optimal parameters estimated by ∂a∂i, and then from this likelihood and the number of parameters of the model calculated the AIC (Akaike 1974). For the constant population size model, there are no optimized parameters, so the AIC is simply minus 2 times the log-likelihood of the model. Model selection was performed for each data set simulated with constant population size, with or without selection. For each model with variable population size, the python script we used to perform parameter optimization and obtain the likelihood of the optimal parameterization has been deposited on GitHub (https://github.com/kern-lab/demogPosSelDadiScripts), as has the script used to obtain the likelihood under the constant population size model.

To perform formal model selection, we asked for a given simulated data set whether any non-equilibrium model had an AIC at least 50 units lower than that of the equilibrium model. If so, we asked whether any of our three non-equilibrium models had an AIC at least 50 units lower than the other two, in which case we selected that model; otherwise we classified the simulated data set as “ambiguous but non-equilibrium.” If no non-equilibrium model had an AIC at least 50 units lower than the equilibrium model, then we conservatively classified the simulated data set as “equilibrium.” We performed parameter inference and model selection for our recurrent hitchhiking data in the same manner with one exception: when performing model selection in many cases the second optimization step failed when fitting our constant-size populations with selection to non-equilibrium demographic models. We therefore conservatively used parameterizations from the initial coarse optimization to calculate AIC scores for our model selection under recurrent hitchhiking.

### Parameter estimation and model selection using ABC

For each of our three non-equilibrium demographic models, we used ABC to estimate the model parameters in each “observed genome” simulated under the model. To this end, we summarized patterns of variation within each genome by calculating the means and variances of π, the number of segregating sites, Tajima's *D*, 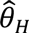 (Fay and Wu 2000), and the number of distinct haplotypes for each of the 500 sampled unlinked loci. We then created a sampling dataset for each demographic model consisting of 5.0×10^5^ simulated genomes evolving in the absence of selection, again with each genome represented by 500 unlinked loci, each of sample size 200. For these sampling datasets, the demographic parameters were drawn uniformly from prior distributions (shown for each model in Table S6), and were summarized by the same set of statistics used for the “observed genomes.” For the bottleneck and contraction-then-growth models, our initial efforts to estimate parameters under neutrality failed to approximate the true parameterizations. We therefore fixed the values of the times of population contraction parameters of these models (referred to as *T_B_* and *T_C_* respectively) during both parameter estimation and model selection: these parameters were always set to the true values during sampling simulations, and their values were not estimated. After this change, we were able to estimate the parameters of each model under neutrality with reasonable accuracy.

We utilized the ABCreg software package (https://github.com/molpopgen/ABCreg) to perform parameter estimation for each of our four demographic models (Thornton 2009). We applied the conventional tangent transformation procedure to the parameters sampled from our prior distributions via passing the –T flag. We also set the tolerance parameter, -t = 0.001, thus retaining 0.1% of our sample data for use in estimating the posterior parameter distributions. ABCreg uses a weighted linear regression approach to the retained sampling simulations in order to estimate the posterior probability densities of the parameters (Beaumont *et al.* 2002). From each parameter’s estimated posterior distribution we used the maximum *a posteriori* estimator (posterior mode) as our point estimate. We also repeated parameter estimation on each observed dataset using the means of summary statistics only.

We performed model selection on our simulated observed genomes with constant population size and varying degrees of positive selection. Our model selection procedure considered each of our four demographic models. For the constant-size model we constructed a new sampling set of 5.0×10^5^ simulated datasets, and for the three variable-size models we used the same sampling sets generated for parameter estimation. We used the R package *abc* to conduct model choice, performing logistic regression-based estimation (using the “mnlogistic” method) of the posterior probabilities of a model (Csilléry *et al.* 2012). For this procedure we set the tolerance parameter to 0.1 for our single-sweep datasets. In the recurrent hitchhiking case we had to increase this parameter to 0.5 in order for the regression procedure to terminate successfully. We separately examined all pair-wise model selection scenarios: constant size vs. growth; constant size vs. contraction-then-growth; constant size vs. bottleneck; bottleneck vs. contraction-then-growth; bottleneck vs. growth demography; and contraction-then-growth vs. growth. We summarized model support with the Bayes factor as calculated by the *abc* package; in cases where this calculation resulted in division by zero (i.e. where the posterior probability of a model was estimated to be zero), we set the Bayes factor to be equal to the largest one observed among datasets with the same combination of demographic model, *f*, and *c*/*s*.

We performed model selection in a manner similar to our approach with ∂a∂i, asking for a given simulated data set whether any non-equilibrium model had a Bayes factor of at least 20 when compared to the equilibrium model. If so, we asked whether any of our three non-equilibrium models had a Bayes factor ≥20 when compared to each of the two, in which we selected that model; otherwise we classified the simulated data set as “ambiguous but non-equilibrium.” Analogous to our model selection with ∂a∂i, if no non-equilibrium model had a Bayes factor ≥20 when compared to the equilibrium model, then we conservatively classified the simulated data set as “equilibrium.” This was done for both our single-sweep and recurrent hitchhiking scenarios.

### Inferring population size histories with PSMC

We ran PSMC in order to infer the history of population size changes of our 15 Mb simulations from which two individuals were sampled (see above). Briefly, we converted our simulation output to the same format generated by running msHOT-lite (https://github.com/lh3/foreign/tree/master/msHOT-lite) with the –l flag. We then ran PSMC’s ms2psmcfa.pl script with default parameters to generate input for PSMC, which we ran with the following parameters: -N25 -t15 -r5 -p ”4+25*2+4+6”. Finally, we ran PSMC’s psmc2history.pl script with default parameters to output the inferred population size history. We then rescaled the output from units of *Ne* to generations and numbers of individuals using the estimated value of *θ*. For each selective scenario, we ran PSMC separately on all 100 simulated population samples. Finally, for the purposes of visualization we obtained a median estimate of population size across time by examining a large number of time points (one every 100 years) across the entire period examined, and at each time point taking the median population size estimate from all 100 simulations.

## Acknowledgements

We thank Jody Hey and Matthew Hahn for feedback on the manuscript. D.R.S. was partially supported by the National Institutes of Health under Ruth L. Kirschstein National Research Service Award F32 GM105231. A.D.K. was supported in part by NIH award no. R01GM078204.

## SUPPLEMENTARY FIGURE AND TABLE LEGENDS

**Figure S1.** Bottleneck model parameter estimates from ∂a∂i, ABC using summary statistic means, and ABC using both means and variances. Parameter estimation was performed on simulated data sets either evolving neutrally, or with some fraction (*f*) of loci used for inference linked to a selective sweep at some distance (measured by *c*/*s*). Each box plot summarizes estimates from 100 replicates for each scenario. Note that *T_B_*, the bottleneck onset time, is absent from this figure because it was fixed to the true value (Methods).

**Figure S2.** Bottleneck model parameter estimates from ∂a∂i, ABC using summary statistic means, and ABC using both means and variances. Parameter estimation was performed on simulated data sets either evolving neutrally, or with some fraction of regions (*f*) used for inference experiencing recurrent hitchhiking at linked loci. Each box plot summarizes estimates from 100 replicates for each scenario. Note that *T_B_*, the bottleneck onset time, is absent from this figure because it was fixed to the true value (Methods). For this analysis ∂a∂i often failed to optimize on runs with a larger fraction of loci experiencing selection, so for ∂a∂i we omitted all datasets with an *f* of 0.7 or greater.

**Figure S3.** Growth model parameter estimates from ∂a∂i, ABC using summary statistic means, and ABC using both means and variances. Parameter estimation was performed on simulated data sets either evolving neutrally, or with some fraction (*f*) of loci used for inference linked to a selective sweep at some distance (measured by *c*/*s*). Each box plot summarizes estimates from 100 replicates for each scenario. Note that *T_B_*, the bottleneck onset time, is absent from this figure because it was fixed to the true value (Methods).

**Figure S4.** Growth model parameter estimates from ∂a∂i, ABC using summary statistic means, and ABC using both means and variances. Parameter estimation was performed on simulated data sets either evolving neutrally, or with some fraction of regions (*f*) used for inference experiencing recurrent hitchhiking at linked loci. Each box plot summarizes estimates from 100 replicates for each scenario.

**Figure S5.** Contraction-then-growth model parameter estimates from ∂a∂i, ABC using summary statistic means, and ABC using both means and variances. Parameter estimation was performed on simulated data sets either evolving neutrally, or with some fraction (*f*) of loci used for inference linked to a selective sweep at some distance (measured by *c*/*s*). Each box plot summarizes estimates from 100 replicates for each scenario. Note that *T_C_*, the time of population contraction, is present only for ∂a∂i because for ABC this parameter was fixed to the true value (Methods).

**Figure S6.** Contraction-then-growth model parameter estimates from ∂a∂i, ABC using summary statistic means, and ABC using both means and variances. Parameter estimation was performed on simulated data sets either evolving neutrally, or with some fraction of regions (*f*) used for inference experiencing recurrent hitchhiking at linked loci. Each box plot summarizes estimates from 100 replicates for each scenario. Note that when performing ABC, time of population contraction (*T_C_*), was fixed to the true value and therefore this parameter is only shown for ∂a∂i.

**Figure S7.** Population size histories inferred by PSMC from simulations with recurrent hitchhiking. Inferred size histories for each of 100 replicate simulations are shown as thin gray lines, and the median across all replicates is shown as the thicker line. In all cases the simulated population’s size was constant throughout. (*A*) Population size histories inferred from simulations with a locus-wide rate 2*NλL* of recurrent hitchhiking, where *λ* is the sweep rate per bp and *L* is the locus size (15 Mb), of 0.01. (*B*) Inferences where 2*NλL* = 0.1. (*C*) 2*NλL* = 1.0. (*D*) 2*NλL* = 10. (*E*) 2*NλL* = 100. (*F*) 2*NλL* = 1000.

**Figure S8.** Population size histories inferred by PSMC from simulations under the contraction-then-growth model. Inferred size histories for each of 100 replicate simulations are shown as thin gray lines, and the median across all replicates is shown as the thicker line. In all cases the simulated population experienced the contraction-then-growth demographic history. (*A*) Population size histories inferred from neutral simulations. (*B*) Inferences from simulations with one selective sweep, for which the fixation time is shown as a dashed green vertical line. (*C*) Inferences from simulations with three recurrent selective sweeps. Fixation times for the two older sweeps are shown as dashed green vertical lines, while the most recent sweep fixed immediately prior to sampling. (*D*) Five recurrent selective sweeps, with fixation times for the four oldest shown as dashed vertical lines; again, the most recent sweep fixed immediately prior to sampling.

**Figure S9.** Differences in AIC between equilibrium and non-equilibrium models when fitted by ∂a∂i to simulated constant-size populations with varying degrees of positive selection. (*A*) AIC scores from simulations experiencing recent selective sweeps. (*B*) AIC scores from simulations experiencing recurrent hitchhiking. For the growth model in panel *A*, a small number of simulated optimized very poorly, leading to large AICs, and therefore large differences between the growth and equilibrium AIC. The upper limit of the *y*-axis of this plot was truncated to allow visualization of AIC differences for the bulk of the data for which optimization was more successful (though box and whisker lengths still reflect the presence of these outliers in the set).

**Figure S10.** Bayes factors from ABC’s model selection comparing equilibrium and non-equilibrium demographic models. (*A*) Bayes Factors from simulations experiencing recent selective sweeps. (*B*) Bayes Factors from simulations experiencing recurrent hitchhiking. Bayes Factors were capped at 10^8^ and 10^-8^ for visualization.

**Table S1.** The fraction of simulated data sets for which each demographic model was selected by ABC when including variances of summary statistics.

**Table S2.** The fraction of simulated data sets with recurrent hitchhiking for which each demographic model was selected by ∂a∂i.

**Table S3.** The fraction of simulated data sets with recurrent hitchhiking for which each demographic model was selected by ABC.

**Table S4.** The fraction of simulated data sets with recurrent hitchhiking for which each demographic model was selected by ABC when including variances of summary statistics.

**Table S5.** Example command lines to simulate each demographic model using discoal.

**Table S6.** Priors on parameter values of each demographic model for ABC sampling simulations.

